# Interplay Between Protein–RNA Binding and Phase Separation Drives Emergent Behavior in RNP Condensates

**DOI:** 10.64898/2026.06.25.734509

**Authors:** Matteo Boccalini, Davide Erba, Matteo Paloni, Alessandro Barducci

## Abstract

Protein–RNA binding and biomolecular condensation are two key processes underlying the assembly and function of ribonucleoprotein (RNP) condensates. However, the understanding of the physical consequences of their interplay is still incomplete. To investigate this coupling, here we develop a minimal coarse-grained molecular model that combines specific, saturable protein–RNA binding with multivalent protein–protein interactions. Our results show that RNA acts as a molecular scaffold whose ability to promote condensation depends on the distribution of bound proteins across RNA molecules. This provides a simple microscopic explanation for both RNA-length-dependent condensation and re-entrant phase behavior, showing that condensate dissolution at high RNA concentration can emerge from entropic effects without requiring explicit electrostatic interactions. Conversely, condensate assembly markedly enhances effective protein–RNA binding, demonstrating that substantial changes in binding behavior can emerge without changes in intrinsic affinity. This provides a general physical mechanism through which condensates can reshape molecular competition between RNA-binding proteins. Together, these findings establish a framework linking RNA binding and biomolecular condensation, illustrating how their interplay governs condensate assembly.

## Introduction

Ribonucleoprotein (RNP) granules are dynamic, membraneless organelles composed of RNA and RNAbinding proteins (RBPs) that play crucial roles in RNA metabolism including RNA processing, regulation of translation, and the storage and degradation of messenger RNAs (mRNAs) Decker and Parker (2012); Ripin and Parker (2023); Otis and Mowry (2023); Pei et al. (2025). RNP granules are ubiquitously distributed throughout the cell, encompassing cytoplasmic assemblies such as processing bodies (P-bodies) and stress granules, as well as nuclear condensates including the nucleolus, nuclear speckles, Cajal bodies, and paraspeckles Decker and Parker (2012); Mao et al. (2011). *In vitro* and *in vivo* studies suggest that RNP granules are formed through a dynamic network of multivalent and heterogeneous interactions between RBPs and RNA Alberti et al. (2019). These interactions can give rise to highly organized and complex condensates Ripin and Parker (2023) which influence spatial organization and RNA localization, thereby generating distinct microenvironments that modulate biochemical reactions Banani et al. (2017).

Despite significant progress, the molecular mechanisms regulating RNP granules remain only partially understood Putnam et al. (2023); Ripin and Parker (2023). RBPs are markedly enriched in intrinsically disordered regions (IDRs) relative to the global proteome Youn et al. (2019); Zhao et al. (2021) and these regions have been widely proposed to promote phase separation through weak intermolecular interactions Molliex et al. (2015); Alberti and Dormann (2019).

Molecular simulations spanning atomistic Paloni et al. (2021); Welsh et al. (2022); Unarta et al. (2024); Boccalini et al. (2025) to coarse-grained (CG) resolution Joseph et al. (2021); Nguyen et al. (2022); Valdes-Garcia et al. (2023); Yasuda et al. (2025) have provided important insights into the molecular interactions governing the behavior of model protein–RNA condensates. Although IDRs alone are often sufficient to drive condensate formation Martin et al. (2020), RBPs also harbor structured RNA-binding domains that mediate specific interactions with RNA Corley et al. (2020). Such interactions are intrinsic to the assembly of RNP systems and have been shown to play central roles in condensate formation and regulation Ripin and Parker (2023). However, a microscopic understanding of how specific protein–RNA interactions and multivalent protein–protein interactions jointly determine condensate behavior remains incomplete. To address this question, we developed a minimal coarse-grained molecular model that combines multivalent protein–protein interactions with specific, saturable protein–RNA binding. By retaining only the essential physical ingredients of the system, the model makes it possible to probe the length and time scales relevant for RNP assemblies. This approach allows us to investigate how specific protein–RNA interactions influence condensate assembly and how the resulting condensates affect the organization of protein–RNA complexes.

## Results

### A Toy Model for RNP formation

We developed an ultra–CG model to capture the physical basis of RNA–protein condensate formation. The central premise is to reproduce the complex interplay between homotypic protein–protein interactions and heterotypic protein–RNA interactions, which together govern phase separation in RNP systems Jankowsky and Harris (2015).

In the model, proteins are represented as patchy particles composed of two distinct sites: a binding site representing the specific domain responsible for RNA recognition and a core site that accounts for the other protein domains, including IDRs, mediating non-specific protein–protein interactions (Fig. 1A). RNA molecules are modeled as flexible linear polymers consisting of interacting beads, each corresponding to a minimal RNA segment capable of engaging a protein binding site (Fig. 1B).

**Figure 1.**
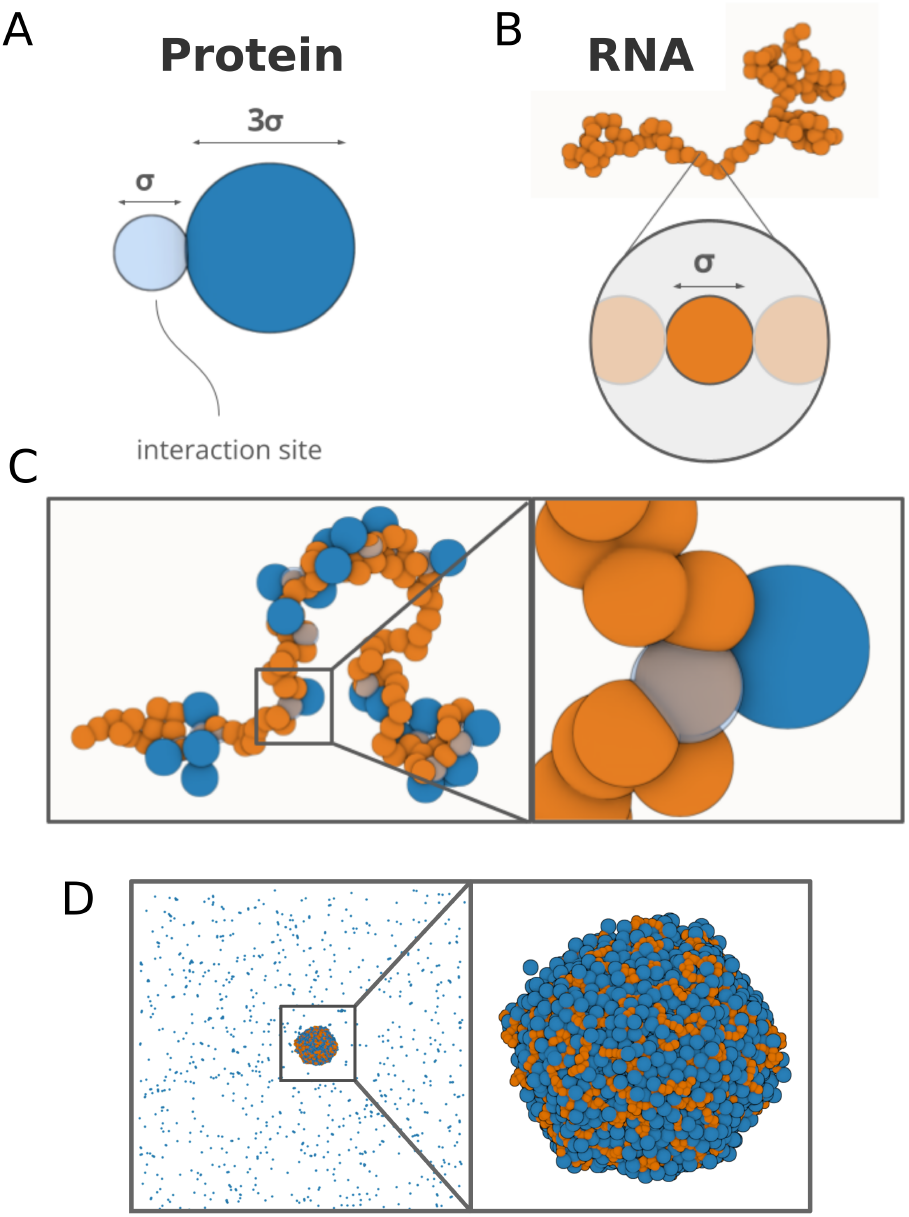
Ultra coarse-grained model. (A) Schematic representation of the protein, modeled as a patchy particle with a specific interaction site capable of binding RNA. (B) RNA chain, where each bead represents an active site able to interact with an individual protein interaction site at a time. (C) Snapshot illustrating how the protein can saturate individual RNA interaction sites with a 1:1 binding stoichiometry. (D) Snapshot of the protein–RNA condensate formed during a simulation.

To capture the selective and saturable nature of protein–RNA binding mediated by RBDs, we implement a short-range attractive interaction between the protein binding site and individual RNA beads, following a strategy inspired by previous work Zhang et al. (2021).

This interaction enforces an effective 1:1 stoichiometry by preventing multiple proteins from simultaneously engaging the same RNA site (Fig. 1C). Conversely, we model homotypic protein–protein interactions, which are assumed to arise from non-specific multivalent contacts between low-complexity disordered regions, using a simple Lennard-Jones like potential Ashbaugh and Hatch (2008) between protein cores.

This simple model successfully reproduces in vitro observations indicating that RNA can promote condensate assembly under conditions where proteins do not phase separate through homotypic interactions Zhang et al. (2015); Molliex et al. (2015); Saha et al. (2016); Hondele et al. (2019); Yang et al. (2020). Under conditions where homotypic protein–protein interactions alone are insufficient to drive condensation, the addition of one equivalent of RNA (corresponding to a ratio of 1 RNA per 100 protein molecules), immediately triggers demixing of the system, despite the protein concentration being far below the saturation threshold observed in the protein-only system (Fig. S2).

### Parameter Space Exploration of Protein–RNA Phase Separation

The model enables independent control of protein–protein and protein–RNA interaction strengths through the parameters *ε*_PP_ and *ε*_PR_, respectively, allowing us to systematically investigate how these two energetic scales individually contribute to macromolecular assembly.

Keeping the total concentrations of protein and RNA constant (as detailed in the simulation protocol section of the Methods), we systematically varied both *ε*_PP_ and *ε*_PR_ (Fig. 2A). The parameter *ε*_PR_ represents the protein–RNA binding affinity and was explored in the range 11–17 k_B_T, corresponding approximately to affinities spanning the millimolar to nanomolar regime (Fig. S3). The protein–protein interaction strength (*ε*_PP_) was varied between 0 and 2.2 k_B_T. Above this value, homotypic protein–protein interactions alone are sufficient to drive phase separation, even in the absence of RNA.

**Figure 2.**
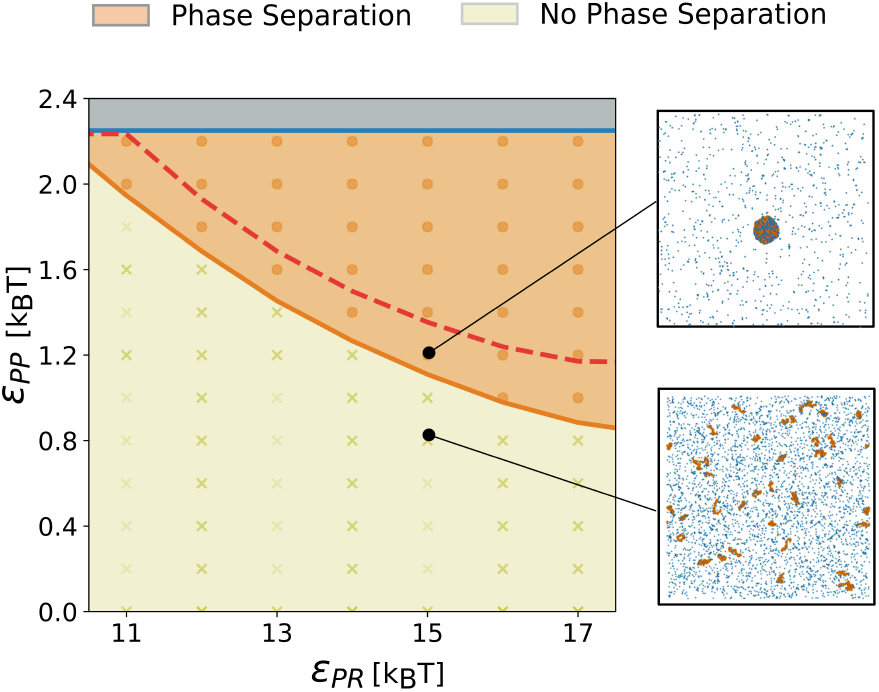
Phase diagram as a function of the parameters *ε*_PR_ and *ε*_PP_. Each point represents a simulation. The orange region indicates where the condensate forms, while the blue region corresponds to values for which the protein undergoes phase separation independently of RNA. The red line represents the phase boundary for systems containing shorter RNA chains (RNA_10_ + proteins). Representative snapshots for condensate-forming and non-condensate-forming conditions are shown on the right.

The resulting phase diagram reveals a broad region of the parameter space (highlighted in orange in Fig. 2A) in which the protein–RNA mixture undergoes phase separation. The shape of the phase boundary indicates that the two interactions contribute cooperatively to condensate formation. For strong protein–protein interactions (e.g., *ε*_PP_ = 2.0 k_B_T), even moderate protein–RNA affinity (*ε*_PR_ = 11 k_B_T) is sufficient to drive condensate formation. Conversely, at lower *ε*_PP_ values (e.g., 1.0 – 1.2 k_B_T), phase separation requires stronger protein–RNA interactions, and therefore higher *ε*_PR_. Notably, RNA-mediated interactions enable phase separation in regimes that would not be accessible even in the presence of very strong protein–protein interactions alone, as showed in Fig. 2 and Fig. S4.

These results show that the model captures the central role of RNA in promoting phase separation and highlight how condensate formation is regulated by the interplay between the intrinsic phase-separation propensity of the protein and the strength of protein–RNA interactions. This behavior can be rationalized by considering a scaffolding role of RNA. In this scenario, RNA molecules confine the proteins bound to them within a restricted spatial region, thereby facilitating protein–protein contacts and promoting condensation.

If RNA acts as a scaffold, its ability to promote phase separation should depend on its length. To test this hypothesis, we “fragmented” the RNA molecules used in our simulations. Specifically, each RNA molecule of length 100 (RNA_100_) was replaced by ten RNA molecules of length 10 (RNA_10_), while keeping the total RNA mass in the system constant. The resulting phase diagram (red curve in Fig. 2A) shows a contraction of the phase-separated region. In other words, under conditions close to the phase boundary, shortening the RNA length suppresses condensate formation (Fig. S4). Consistent with this observation, experimental studies have reported that RNA length modulates phase separation behavior Saha et al. (2016); Garcia-Jove Navarro et al. (2019); Yang et al. (2020); Roden and Gladfelter (2021).

These results support a model in which RNA acts as a scaffold that promotes the local enrichment of proteins by physically constraining them within a finite volume. In this scenario, the diffusion of the RNA chain effectively replaces that of individual proteins, resulting in a reduction of translational entropy. This entropic penalty is compensated by the enthalpic gain arising from multivalent interactions and the resulting phase separation.

### Entropy-Driven Re-entrant Phase Separation

We next asked whether our model can reproduce the re-entrant behavior observed experimentally, in which RNA can both promote and dissolve biomolecular condensates. Previous *in vitro* studies have shown that RNA not only enhances phase separation but, at sufficiently high concentrations, can also destabilize and dissolve condensates Banerjee et al. (2017); Maharana et al. (2018); Alshareedah et al. (2019); Wadsworth et al. (2024).

To test this prediction within our framework, we titrated the system by progressively increasing RNA concentration while keeping the protein concentration constant. The results are shown in Fig. 3A, where concentrations are expressed as the ratio of RNA beads to protein. For each condition, we quantified the fraction of protein partitioning into the condensed phase (Fig. 3B).

**Figure 3.**
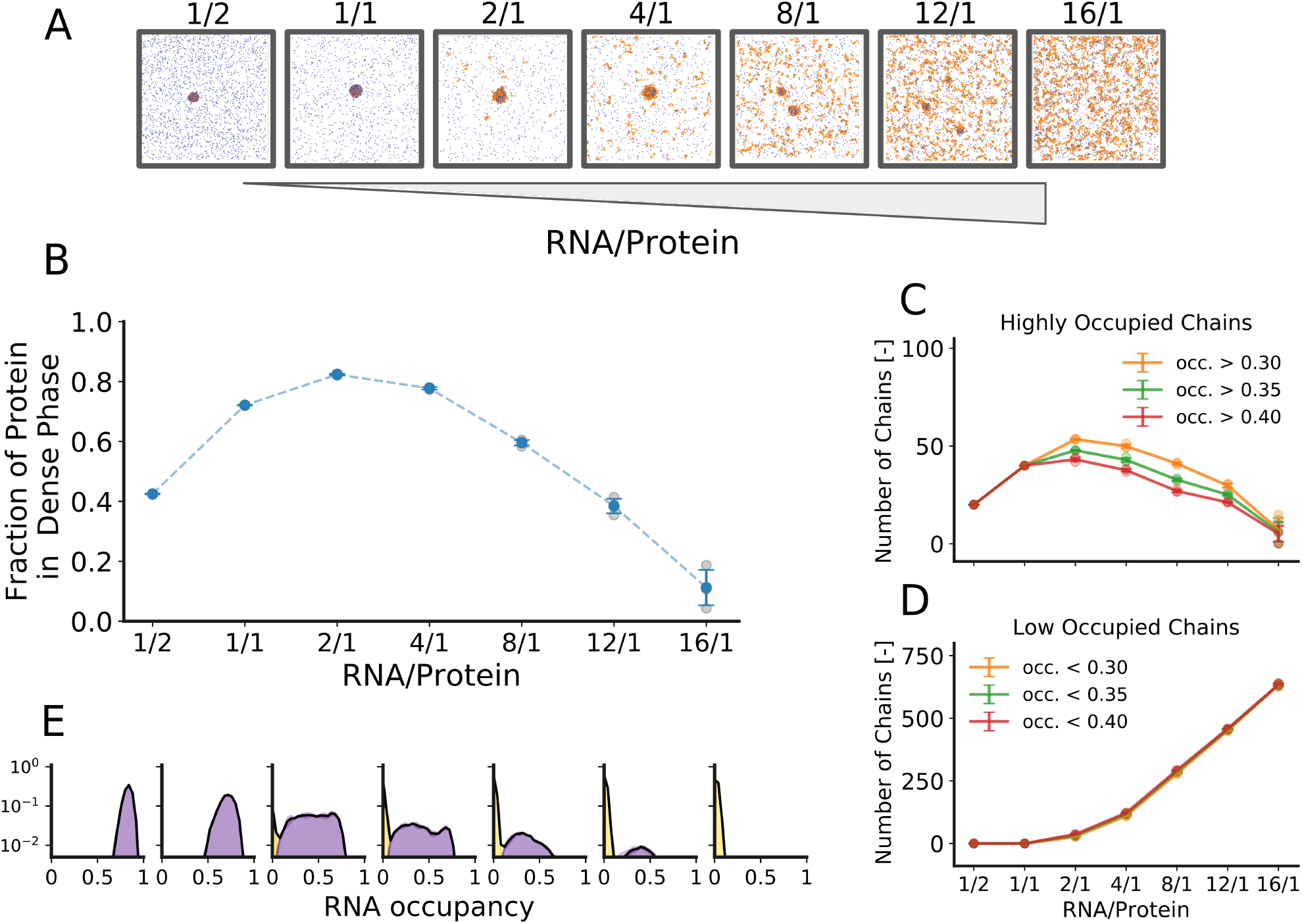
Re-entrant phase diagram. (A) System titration by varying RNA concentration while keeping protein constant. Insets show snapshots of the simulation box, with the bottom panel reporting the percentage of protein in the condensed phase. (B) Fraction of protein in the condensed phase as a function of the RNA/protein ratio. (C) Number of RNA chains exhibiting protein occupancy greater than 30%, 35%, and 40% as a function of the RNA/protein ratio. (D) Number of RNA chains exhibiting protein occupancy lower than 30%, 35%, and 40% as a function of the RNA/protein ratio. (E) Distribution of RNA chain occupancy across the RNA/protein ratio. Purple indicates the fraction of chains within the dense phase, and yellow corresponds to chains in the dilute phase.

The introduction of small amounts of RNA was sufficient to trigger condensate formation. At an RNA:protein ratio of 0.5:1, approximately 40% of the protein partitioned into the dense phase, closely matching the amount required to saturate the available RNA-binding sites. Increasing the RNA concentration to 1:1 further enhanced condensation, with 70% of the protein residing in the dense phase. Condensation peaked at 80 % at an RNA:protein ratio of 2:1. At this point, however, a fraction of RNA chains began to dissociate from the condensate. Further increases in RNA concentration led to a progressive decline in the condensed protein fraction. At an extreme ratio of 16:1, condensate formation was strongly suppressed, with the dense phase nearly completely dissolved.

To understand the origin of this non-monotonic behavior, we revisited the hypothesis that RNA acts as a molecular scaffold, locally constraining proteins and stabilizing their interactions. Within this framework, the effectiveness of RNA as a scaffold depends on its ability to recruit multiple proteins simultaneously. We therefore quantified the occupancy of RNA chains, defined as the fraction of protein-binding sites engaged along each RNA molecule. At low and intermediate RNA concentrations, the system contains a substantial population of highly occupied RNA chains, consistent with efficient multivalent scaffolding. As RNA concentration increases, however, the occupancy distribution progressively shifts toward sparsely occupied RNA molecules, while the population of highly occupied chains decreases markedly in the re-entrant regime (Fig. 3C-3D). These observations suggest that condensate stability is associated not simply with the total number of RNA-bound proteins, but with the presence of sufficiently populated RNA scaffolds capable of locally concentrating proteins. At high RNA concentration, proteins can distribute across a large number of RNA molecules rather than accumulating on a smaller subset of highly occupied chains. This redistribution reduces the local protein concentration along individual RNA scaffolds and weakens the multivalent interactions required to stabilize the condensate. In this regime, dispersing proteins over many RNA molecules becomes entropically favorable relative to concentrating them onto a smaller number of highly occupied scaffolds and the resulting loss of local multivalent stabilization ultimately drives condensate dissolution.

A more detailed analysis of the occupancy distributions (Fig. 3E) reveals that, under conditions where both condensed and dilute phases coexist, RNA molecules separate into distinct occupancy subpopulations. Interestingly, under these conditions RNA molecules are selectively partitioned according to their occupancy state. Highly occupied RNA chains are preferentially partitioned into the condensed phase, whereas sparsely occupied RNA molecules remain predominantly in the dilute phase (Fig. 3E and Fig. S5).

Together, these results suggest that RNA occupancy and condensate formation are tightly coupled. RNA molecules act as scaffolds whose ability to promote condensation depends on their capacity to locally concentrate proteins. At the same time, the higher occupancy of RNA chains within condensates suggests that phase separation itself may reshape the effective binding landscape of RNA–protein interactions.

### Phase Separation Enhances Protein–RNA Affinity

We therefore next asked whether condensate formation could, in turn, modulate the effective affinity of proteins for RNA. To address this, we employed our CG model to systematically investigate the effect of phase separation on protein-RNA affinity. We kept the protein-RNA interaction strength constant at a micromolar-range affinity (*ε*_PR_ = 15 k_B_T), while progressively increasing the homotypic protein-protein interaction strength (*ε*_PP_), in an 1:1 protein:RNA system.

The extent of binding was quantified via the fraction of total protein molecules bound to RNA. As shown in Fig. 4A, in the absence of protein-protein interactions (*ε*_PP_ = 0 k_B_T), approximately 10% of the protein population is RNA-bound. Gradual increases in *ε*_PP_ led to modest increases in binding, which remained below 20%. However, beyond a critical *ε*_PP_ threshold (*ε*_PP_ > 1.0 k_B_T), phase separation was observed, and the fraction of RNA-bound protein sharply increased, approaching full RNA occupancy at high *ε*_PP_ values. These results indicate that condensates act as a strong proteinRNA binding enhancer.

**Figure 4.**
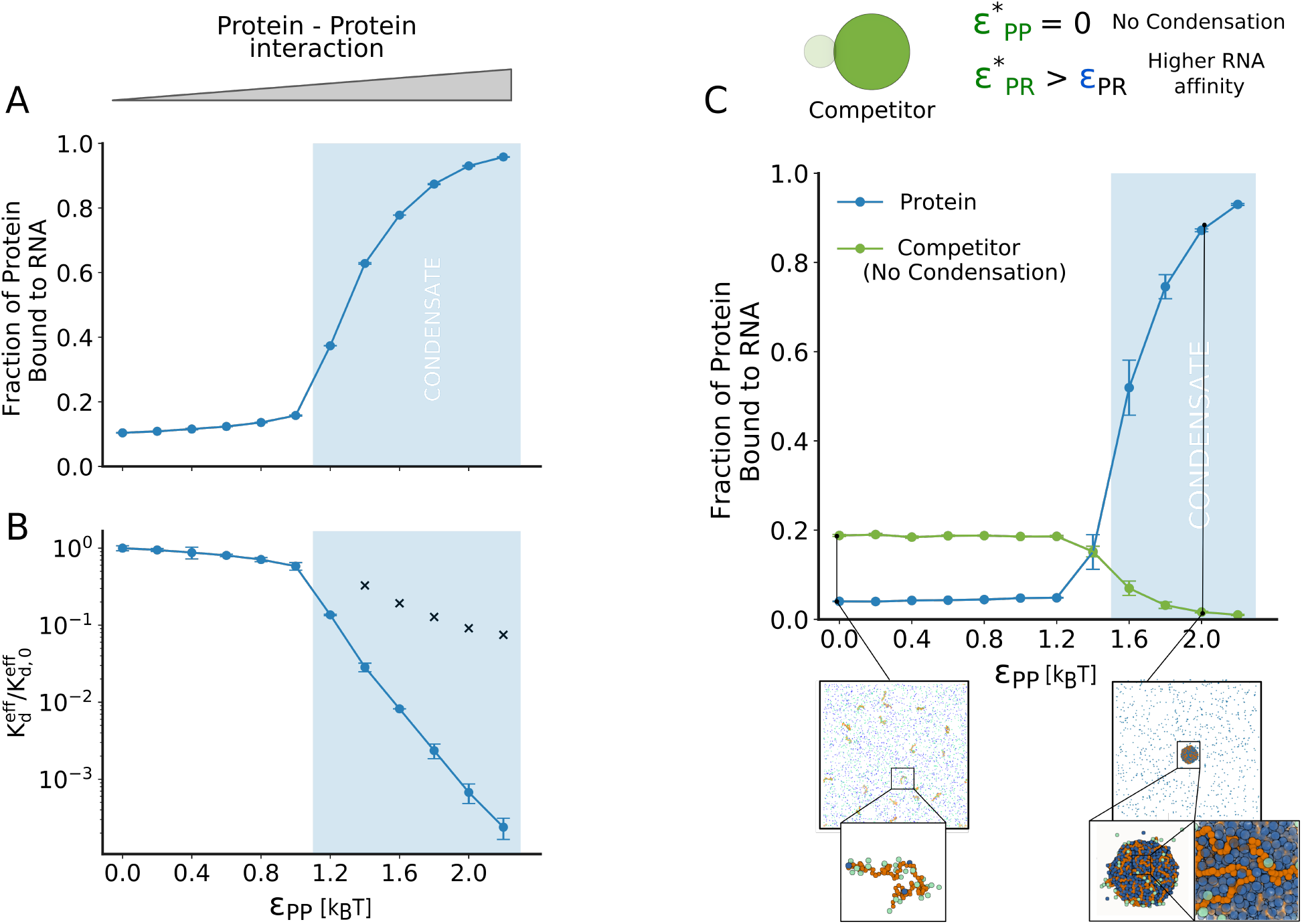
Condensates enhance protein-RNA binding. (A) Fraction of protein bound to RNA as a function of protein-protein interaction strength (*ε*_PP_). The blue region indicates values of *ε*_PP_ where condensates form. (B) Effective 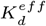 of the system as a function of *ε*_PP_, normalized by the 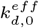 obtained without protein-protein interaction (*ε*_*PP*_ = 0). The blue region highlights *ε*_PP_ values corresponding to condensate formation, where the protein-RNA affinity increases up to 4-fold. Black crosses indicate dissociation constant evaluated using only concentrations within the condensed phase (see SI methods). (C) Fraction of protein bound to RNA as a function of *ε*_PP_; blue corresponds to the protein and green to the protein competitor. The blue region indicates conditions where the protein undergoes phase separation. In the upper part, schematic representation of the competitor protein, which exhibits higher affinity for RNA (larger *ε*_PR_ than the protein) but lacks protein-protein interactions 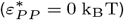. In the bottom part, snapshots of the system at *ε*_*PP*_ = 0 k_B_T and *ε*_*PP*_ = 2.0 k_B_T.

To better quantify this effect, we computed an effective dissociation constant 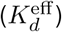 across the entire system, independent of whether molecules are located in the dilute or condensed phase. As shown in Fig. 4B, 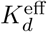 remains nearly constant prior to phase separation, consistent with the weak cooperative effects observed at low *ε*_PP_ values. Once the system enters the phase-separated regime, however, 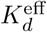 decreases dramatically by two to three orders of magnitude, corresponding to a strong enhancement in effective protein–RNA affinity.

Importantly, this enhancement arises predominantly from the localization generated by phase separation itself. When the dissociation constant is evaluated considering only concentrations within the condensed phase (crosses in Fig. 4B and Fig. S7), the reduction in *K*_*d*_ is substantially smaller. This observation suggests that condensate formation enhances effective protein–RNA binding primarily through local enrichment effects, rather than through a major change in intrinsic affinity within the dense phase.

To test whether the localization-driven enhancement of effective affinity can alter molecular competition, we designed an *in silico* competition experiment. Specifically, we extended our coarse-grained model to include a second protein species, termed the competitor, which can also bind RNA but lacks favorable homotypic interactions and therefore does not phase separate under these conditions.

Structurally, the competitor is identical to the original protein except that homotypic protein–protein intercations are removed 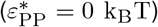. To establish a direct competition for RNA binding, the competitor was assigned a stronger protein–RNA interaction strength 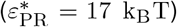 than the original protein (*ε*_PR_ = 15 k_B_T). A schematic of the system is shown in Fig. 4C. We simulated a ternary mixture composed of equal amounts of protein and competitor together with RNA (0.5:0.5:1 protein:competitor:RNA).

As previously done, we varied the self-interaction strength of the original protein (*ε*_PP_) to assess its effect on competitive RNA binding. When *ε*_PP_ = 0 k_B_T, the competitor dominates RNA binding owing to its stronger intrinsic affinity: approximately 20% of competitor molecules are RNA-bound, whereas binding of the original protein remains marginal (Fig. 4C). Moderate increases in *ε*_PP_ produce only minor changes, and the competitor continues to outcompete the protein.

However, once the system enters the phase-separated regime, a sharp redistribution of RNA binding occurs. The local enrichment generated by condensate formation strongly favors recruitment of the condensate-forming protein, which becomes the dominant RNA-bound species despite its weaker intrinsic affinity. Concomitantly, RNA binding by the competitor decreases dramatically, approaching zero at high *ε*_PP_ values (Fig. 4C). These results demonstrate that phase separation can override intrinsic binding preferences through collective localization effects, effectively reshaping molecular competition for RNA.

## Discussion

Protein–RNA binding and biomolecular condensation are fundamental processes underlying the assembly and function of ribonucleoprotein condensates. Using a minimal molecular model, we show that their coupling is sufficient to generate a range of complex behaviors with potential functional relevance. Rather than acting independently, RNA binding and phase separation become reciprocally linked processes, with each reshaping the other and jointly determining condensate assembly and organization.

Consistent with previous experimental and theoretical studies Roden and Gladfelter (2021); Sanchez-Burgos et al. (2022); Wadsworth et al. (2024), our results support a picture in which RNA chains act as molecular scaffolds that promote condensation by bringing multiple bound proteins into close proximity. However, our occupancy analysis suggests that the effects of RNA length and concentration can be rationalized in terms of a common variable: the population of highly occupied RNA chains that serve as molecular scaffolds for condensation.

This perspective provides a mechanistic explanation for re-entrant phase behavior, a phenomenon widely observed in vitro Maharana et al. (2018); Banerjee et al. (2017) and increasingly viewed as a potential mechanism for the dynamic regulation of condensates involved in gene expression and RNA metabolism in vivo Henninger et al. (2021); Muzzopappa and Erdel (2025). Within this framework, increasing RNA beyond an optimal concentration progressively redistributes proteins across a larger number of RNA molecules, thereby reducing the population of highly occupied RNA scaffolds that stabilize the condensed phase. Re-entrant dissolution occurs when this population becomes insufficient to sustain condensation. From a thermodynamic perspective, this transition reflects the balance between the enthalpic stabilization provided by the condensed phase and the entropic gain associated with distributing proteins across many RNA molecules. More broadly, our results suggest that re-entrant phase behavior can arise from competition between condensation and alternative protein–RNA organizations that become increasingly favored at high RNA concentrations, a view that is qualitatively consistent with recent experimental evidence for competition between phase separation and protein–RNA clustering Makasewicz et al. (2026). While re-entrant phase behavior is frequently rationalized in terms of electrostatic mechanisms, such as charge inversion Banerjee et al. (2017); Alshareedah et al. (2019), our results show that explicit electrostatic interactions are not strictly required for its emergence, potentially providing an alternative explanation for reentrant behavior observed in weakly charged systems Zhang et al. (2015); Fatti et al. (2025).

Protein–RNA interactions play central roles in numerous cellular processes, including RNA metabolism, localization, and post-transcriptional regulation Gerstberger et al. (2014); Gebauer et al. (2021). Because ribonucleoprotein condensates are increasingly implicated in the regulation of these processes, understanding how condensation influences protein–RNA binding is of fundamental importance. While protein–RNA interactions are widely recognized as key determinants of condensate assembly, our results demonstrate that the relationship is reciprocal. Condensate formation strongly enhances protein–RNA binding. Analysis of the binding equilibria suggests that this enhancement arises primarily from the enrichment of proteins and RNA within the condensed phase, demonstrating that substantial changes in binding behavior can emerge even without major changes in intrinsic affinity. More generally, coupling protein–RNA binding to an inherently cooperative process, such as condensate assembly, may amplify relatively modest changes in the factors controlling condensation into substantial changes in binding behavior. Furthermore, this enhancement of protein–RNA binding through localization may have important implications for molecular selectivity. In particular, our results suggest that the outcome of molecular competition between RBPs may be determined not only by intrinsic protein–RNA affinity, but also by the propensity of an RNA-binding protein to participate in condensate formation. This finding highlights a general mechanism by which condensates can influence the composition of RNA–protein assemblies, thereby modulating which proteins are preferentially recruited to shared RNA targets.

The present model was intentionally designed to explore the consequences of coupling protein–RNA binding to protein-driven condensation. As a consequence, several factors that may contribute to the behavior of cellular condensates are not explicitly represented, including RNA–RNA interactions, which have recently emerged as potential contributors to condensate organization, as well as sequence-specific molecular recognition, conformational changes, and alterations of the physicochemical environment within condensates Van Treeck and Parker (2018); Roden and Gladfelter (2021). Incorporating such features into the present framework will be important for connecting the general mechanisms identified here to the behavior of specific biological systems. Nevertheless, the emergence of re-entrant phase behavior, enhanced protein–RNA binding, and altered molecular selectivity within such a minimal framework demonstrates that these phenomena do not require these additional layers of complexity. More broadly, our results suggest that occupancy-dependent RNA scaffolding and condensation-enhanced binding represent generic consequences of the reciprocal coupling between RNA binding and condensation, providing a simple framework for understanding how RNP condensates assemble, reorganize, and regulate molecular interactions.

## Methods

To model protein–RNA interactions, we introduce a specific attractive interaction between the protein binding site and individual RNA beads. Our goal is to enforce a 1:1 stoichiometry, where each RNA bead can bind to at most one protein. To achieve this, we use a shortrange attractive potential between the RNA beads and the protein binding site, inspired by approaches previously proposed in the literature Zhang et al. (2021). However, this potential alone does not enforce 1:1 binding. To impose saturable interactions, we introduce a strong repulsive homotypic interaction between protein binding sites, which prevents multiple proteins from simultaneously binding the same RNA bead. Further details about the simulation protocol and analysis methods are available in the SI.

## Acknowledgments

We acknowledge the support of the Swiss National Science Foundation (SNSF) under grant CRSII5_193740 and of the French Agence Nationale de la Recherche (ANR) under grant ANR-21-CE30-0001. M.B. was supported by funds of the LabMuse EpiGenMed.

