## Supplementary Information for "Interplay Between Protein–RNA Binding and Phase Separation Drives Emergent Behavior in RNP Condensates"

### Coarse-grained model

A minimal coarse-grained model was constructed to investigate how the interplay between homotypic protein self-association and heterotypic protein–RNA interactions governs condensate assembly. Proteins were represented as patchy particles composed of two interaction sites: a protein core, representing low-complexity domains that mediate protein self-association, and an RNA-binding site representing a specific RNA-binding domain. RNA molecules were modeled as flexible self-avoiding polymers, in which each monomer represents an RNA segment that can engage a protein binding site.
The connectivity of both proteins and RNA molecules was maintained using harmonic bond potentials,

$$V_{\text{bond}}(r) = \frac{\kappa}{2}(r - r_0)^2,$$

where $\kappa$ is the bond spring constant and $r_0$ is the equilibrium bond length. For consecutive RNA beads, we set $r_0 = 1.5\sigma$, while for protein beads we set $r_0 = 2.3\sigma$, while for both interacting couples $\kappa = 10^3$ kJ mol$^{-1}$ nm$^{-2}$
Protein–RNA interactions were modeled by a short-range attractive potential acting between protein binding sites and RNA beads,

$$V_{PR}(r, r_{PR}) = \begin{cases} -\frac{1}{2}\varepsilon_{PR}\left(1 + \cos\frac{\pi r}{r_{PR}}\right) & r \leq r_{PR} \\ 0 & r > r_{PR} \end{cases}$$

where $\varepsilon_{PR}$ determines the interaction strength and $r_{PR} = \sigma$ defines the interaction range. This potential was adapted from previous coarse-grained models of valence-limited interactions Zhang et al. (2021).
Homotypic protein–protein interactions were modeled using an Ashbaugh–Hatch (AH) potential Ashbaugh and Hatch (2008) acting between protein core beads,

$$V_{PP}(r, r_{PP}) = \begin{cases} \Phi_{LJ}(r, r_{PP}) + (1 - \varepsilon_{PP}) & r \leq 2^{1/6} r_{PP} \\ \varepsilon_{PP}\Phi_{LJ}(r, r_{PP}) & r > 2^{1/6} r_{PP} \end{cases}$$

where $\Phi_{LJ}(r, r_{PP})$ is the Lennard-Jones potential with $r_{PP} = 3\sigma$, and $\varepsilon_{PP}$, expressed in $k_B T$ units, controls the strength of protein self-association.
We adopted the WCA potential Weeks et al. (1971) to prevent the interactions among all the other couples of bead types:

$$V_{\text{rep}}(r, r_0) = \begin{cases} \Phi_{LJ}(r, r_0) + 1, & r \leq 2^{1/6} r_0 \\ 0 & r > 2^{1/6} r_0 \end{cases}$$

where $\Phi_{LJ}(r, r_0)$ is the Lennard-Jones potential computed for a couple $i - j$ such that $r_0$ is the average of their effective sizes. The combination of WCA and $V_{PR}$ enforces valence-limited protein–RNA interactions, since it suppresses the simultaneous association of multiple proteins onto a single RNA bead, resulting in effectively saturable 1:1 binding.

### Coarse-grained model parameters

| Particle type | Mass [Da] | Diameter [$\sigma$] |
|---|---|---|
| RNA bead | 100 | 1 |
| Protein core bead | 100 | 3 |
| Protein site bead | 100 | 1 |

**Table 1.** Particle types parameters.

### Simulation protocol

All simulations were performed using GROMACS 2019 Abraham et al. (2015). First, all the RNA chain and proteins were randomly inserted in a highly concentrated box with `gmx insert-molecules`. Then, condensate formation were enhanced by performing a 4 ns long simulation in the NPT ensemble, with pressure controlled by Parrinello-Rahman barostat Parrinello and Rahman (1981), using an isotropic reference pressure of $2.0$ bar, and temperature

| | RNA bead | Protein core bead | Protein site bead |
|---|---|---|---|
| RNA bead | $V_{rep}$ | $V_{rep}$ | $V_{PR}$ |
| Protein core bead | | $V_{PP}$ | $V_{rep}$ |
| Protein site bead | | | $V_{rep}$ |

**Table 2.** Interaction type for all the particle couples in the system.

(310 K) by a Langevin thermostat. This run allowed to obtain a clear and dense droplet in the centre of the box. Finally, we adjust the final concentration of the system by increasing the box size (we assume $\sigma = 1$ nm). Production simulations were run in the NVT ensemble, where temperature (310 K) was controlled using a Langevin thermostat with a friction coefficient of 0.01 $\mathrm{ps}^{-1}$, and the equations of motion were integrated with a time step of 10 fs.

***Parameter Phase Diagram (Figure 2 and Supplementary Figure 4).*** The phase diagram was constructed by simulating a system containing 4000 protein molecules and 40 RNA chains 100 bead long (RNA100). The corresponding concentrations were 50 $\mu$M protein and 0.5 $\mu$M RNA, resulting in a 1:1 stoichiometric ratio in terms of binding equivalents. For RNA10 (10 beads length; red line in Figure 2), the RNA concentration was set to 5 $\mu$M to preserve the same number of binding equivalents. Phase behavior was characterized by systematically varying the protein-protein interaction strength, $\varepsilon_{PP}$, and the protein-RNA interaction strength, $\varepsilon_{PR}$. Specifically, $\varepsilon_{PP}$ was varied between 0.0 and 2.2 $k_{\mathrm{B}}T$, while $\varepsilon_{PR}$ was varied between 11 and 17 $k_{\mathrm{B}}T$. Each system was initialized from a preformed protein-RNA condensate and simulated for 3 µs. We repeated the same process by using 400 RNA chains 10 bead long (RNA10) to establish the phase diagram coloured with red diagonals in SF.4.

***RNA Titration and Re-entrant Phase Diagram (Figure 3).*** The re-entrant phase diagram was obtained by simulating 4000 proteins at a fixed concentration of 30 $\mu$M while varying the concentration of RNA100 chains. RNA chain concentrations of 0.15, 0.30, 0.60, 1.20, 2.40, 3.60, and 4.80 $\mu$M were considered, corresponding to 20, 40, 80, 160, 320, 480, and 640 RNA100 chains, respectively. For all systems, the interaction parameters were kept constant at $\varepsilon_{PP} = 1.6\,k_{\mathrm{B}}T$ and $\varepsilon_{PR} = 15\,k_{\mathrm{B}}T$. Simulations were initialized from preformed protein-RNA condensates and evolved for 3 $\mu$M. Each condition was independently replicated three times.

***Condensate-enhanced RNA Affinity (Figures 4A-B).*** The simulated system consisted of 4000 proteins and 40 RNA100 chains, corresponding to protein and RNA concentrations of 50 $\mu$M and 0.5 $\mu$M, respectively, maintaining a 1:1 stoichiometric ratio in terms of binding equivalents. Simulations were performed at a fixed protein-RNA interaction strength of $\varepsilon_{PR} = 15\,k_{\mathrm{B}}T$, while $\varepsilon_{PP}$ was varied between 0.0 and 2.2 $k_{\mathrm{B}}T$. Each simulation was run for 3 $\mu$M and repeated in three independent replicates.

***Competition Simulations (Figures 4C).*** Simulations were performed using a system composed of 2000 proteins, 2000 competitor proteins (Protein*), and 40 RNA100 chains, corresponding to concentrations of 25 $\mu$M, 25 $\mu$M, and 0.5 $\mu$M, respectively. The protein-RNA interaction strength was set to $\varepsilon_{PR} = 15\,k_{\mathrm{B}}T$, whereas the competitor protein exhibited a stronger RNA affinity, with $\varepsilon^*_{PR} = 17\,k_{\mathrm{B}}T$. To isolate competitive RNA-binding effects, self-association of the competitor protein was disabled by setting $\varepsilon^*_{PP} = 0$. Simulations were performed for $\varepsilon_{PP}$ values ranging from 0.0 to 2.2 $k_{\mathrm{B}}T$. Each system was initialized from a preformed mixed Protein–Protein*–RNA condensate and evolved for 3 µs. All conditions were simulated in triplicate.

***Protein–RNA Affinity (Figures S3).*** The simulated system consisted of a single protein and a single RNA bead in a box of side 32 nm, corresponding to a concentration of 50$\mu$M. We tested the protein-RNA interaction strength in the range 9-17 $k_{\mathrm{B}}T$, simulated in three independent, $3 \times 20\mu$s-long replicates yielding a total of $3 \times 2 \cdot 10^7$ evenly distributed frames. We estimated the $k_D$ as explained below, with errors obtained by propagating the binding probability.

### Analysis Methods

Analyses were performed using GROMACS Abraham et al. (2015) and MDAnalysis Michaud-Agrawal et al. (2011) functions and modules.

***Clustering and droplet volume.*** The population of condensed droplets were determined by using custom python script taking advantage of the NetworkX library Hagberg et al. (2008). At each frame, a graph of all protein beads was built by adding link among any $i - j$ protein couple whose distance was less than $r_{co} = 6\sigma$.
The droplet, if present, corresponded to the largest connected component of the system and the method adopted allows to monitor its time dissolution in case of low energetic parameters. The droplet volume is estimated from the gyration tensor S computed over all particles belonging to the droplet. The eigenvalues of S, denoted by $\lambda_1$,

$\lambda_2$, and $\lambda_3$, represent the squared principal semi-axes of the corresponding gyration ellipsoid. Under the ellipsoidal approximation, the droplet volume is calculated as

$$V = \frac{4}{3}\pi\sqrt{\lambda_1\lambda_2\lambda_3}.$$

***RNA occupancy.*** RNA occupation was computed in a bead-wise manner using an energetic criterion, by defining as *bound* all RNA-site couples whose distance $\tilde{r}$ is such that

$$V_{PR}(\tilde{r}) \leq -2k_B\mathsf{T} = E_B$$

following Jost Lopez et al. (2020). This corresponds to a $\varepsilon_{PR}$-dependent interaction distance

$$\tilde{r} \leq \frac{1}{\pi}\cdot\arccos\left(\frac{-2\cdot E_B}{\varepsilon_{PR}} - 1\right).$$

***Dissociation constant $K_d$.*** The effective $K_d$ was computed as:

$$K_d^{eff} = \frac{1}{N_{Avo}V}\frac{(1-P)^2}{P}$$

where $P$ si the protein-RNA binding probability computed as the fraction of RNA monomers which are occupied. Its error $\delta K$ was computed by propagating the binding probability error $\delta P$ obtained from independent replicates as:

$$\delta K = \left|\frac{\partial K}{\partial P}\right|\cdot\delta P = \frac{1}{N_{Avo}V}\frac{|1-P|^2}{P^2}\cdot\delta P$$

***Dissociation constant in condensate $K_d^{C.}$.***

$$K_d^{\text{condensate}} = \frac{[R_c][P_c]}{[RP_c]}$$

where $[R_c]$ and $[P_c]$ denote the molar concentrations of unbound RNA (monomers) and proteins inside the condensate, respectively, while $[RP_c]$ denotes the concentration of RNA–protein complexes in the condensate.
We can rewrite the molar concentrations in terms of particle numbers:

$$[P_c] = \frac{p_{\text{tot}} - p_{c,o} - p_{n,o} - p_{n,u}}{V_c}$$

$$[R_c] = \frac{r_{\text{tot}} - r_{c,o} - r_{n,o} - r_{n,u}}{V_c}$$

where $V_c$ is the volume of the condensate. Here $p_{\text{tot}}$ is the total number of proteins in the system, $p_{c,o}$ is the number of proteins bound to RNA monomers inside the condensate, $p_{n,o}$ is the number of proteins bound to RNA monomers outside the condensate, and $p_{n,u}$ is the number of unbound proteins located outside the condensate. The same notation applies to RNA.
In all simulations used to estimate the values reported in Fig. 4B, RNA chains never leave the condensate, so $r_{n,o} = r_{n,u} = 0$. Similarly, RNA–protein complexes outside the condensate are not observed, so $p_{n,o} = 0$. Moreover, each RNA–protein complex contains one RNA monomer and one protein, such that

$$n_{c,o} = p_{c,o} = r_{c,o}.$$

In our simulations, the total number of proteins equals the total number of RNA monomers, and we denote this common value as $n_{\text{tot}}$:

$$n_{\text{tot}} = p_{\text{tot}} = r_{\text{tot}}.$$

With these simplifications, we can rewrite the condensate dissociation constant as

$$K_d^{\text{cond.}} \approx \frac{1}{V_c}\frac{(n_{\text{tot}} - n_{c,o})(n_{\text{tot}} - n_{c,o} - p_{n,u})}{n_{c,o}}.$$

Defining the fraction of bound species as $x = \frac{n_{c,o}}{n_{\text{tot}}}$ and the fraction of unbound proteins in the dilute phase as $\varphi = \frac{p_{n,u}}{n_{\text{tot}}}$, we can rewrite $k_D^{\text{cond.}}$ as

$$K_d^{\text{cond.}} \approx \frac{n_{\text{tot}}}{V_c}\frac{(1-x)(1-x-\varphi)}{x}.$$

Thus, $K_d^{\text{cond.}}$ can be computed from simulations by measuring the bound fraction $x$, the dilute-phase protein fraction $\varphi$, and the condensate volume $V_c$ (Fig. S7).

### Supplementary Information - Figures

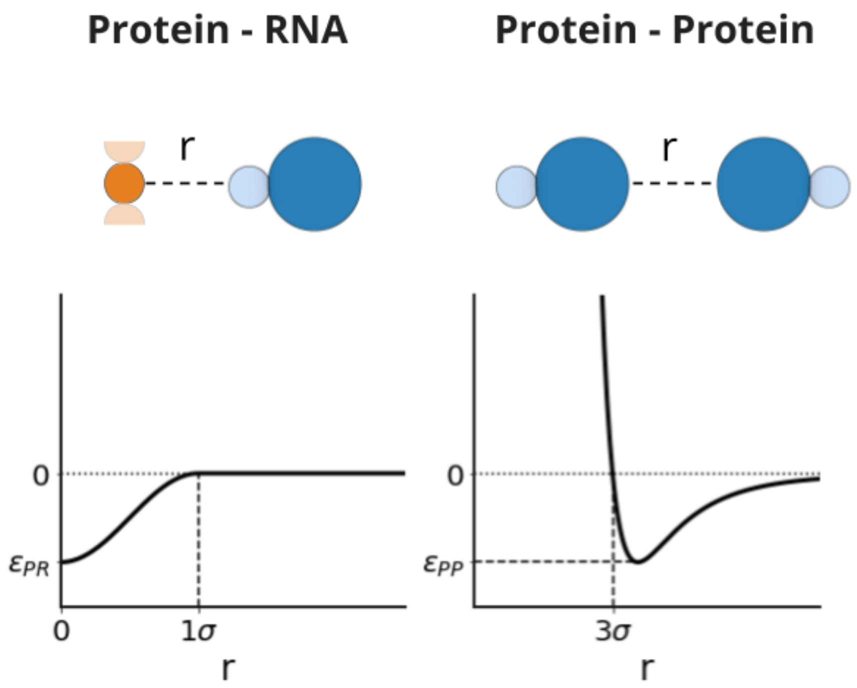


**Figure S1.** Coarse-grained interaction potentials. Left: interaction between an RNA monomer (orange) and a protein binding site (light blue). The corresponding interaction potential, $V_{PR}$, is shown as a function of the interparticle distance. Right: interaction potential between protein core sites (blue). The corresponding potential, $V_{PP}$, is plotted as a function of the interparticle distance.

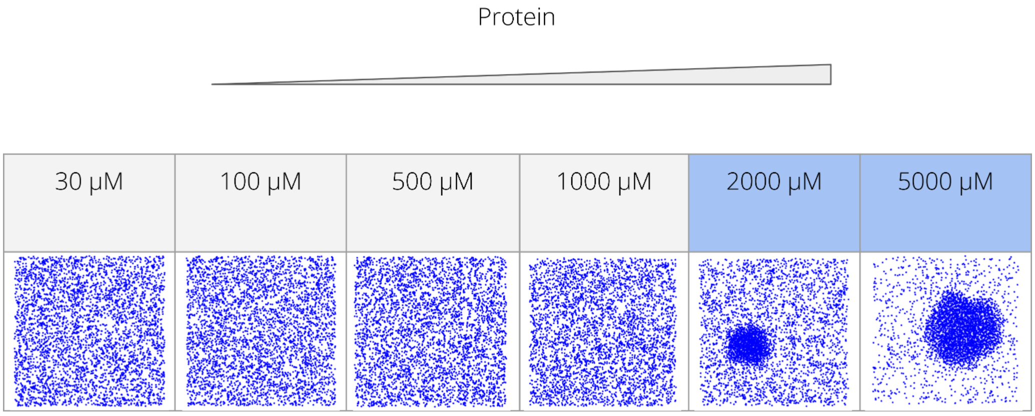


**Figure S2.** Phase diagram as a function of protein concentration. Simulation snapshots are shown for protein concentrations of 30, 100, 500, 1000, 2000, and 5000 $\mu\mathrm{M}$. Condensate formation is observed at protein concentrations equal to or greater than 2000 $\mu\mathrm{M}$. Simulation parameters were $\varepsilon_{PR} = 15\ k_BT$ and $\varepsilon_{PP} = 1.6\ k_BT$.

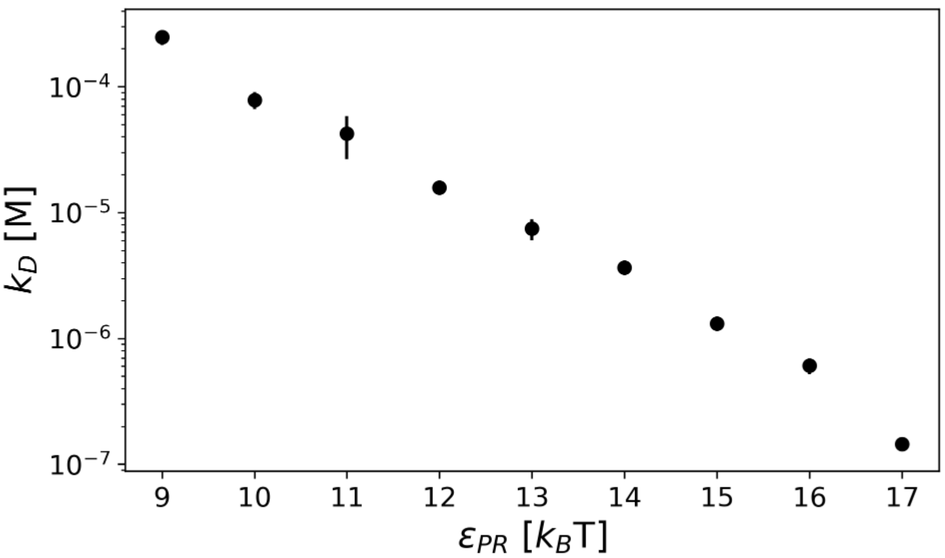


**Figure S3.** Protein-RNA dissociation constant computed by varying $\varepsilon_{PR}$ value for a system made by an RNA monomer and a single protein at a concentration of 50 M. Errors showed are computed by propagating the error of the bound population obtained from three independent replicates on the effective $K_d$ function described in the methods.

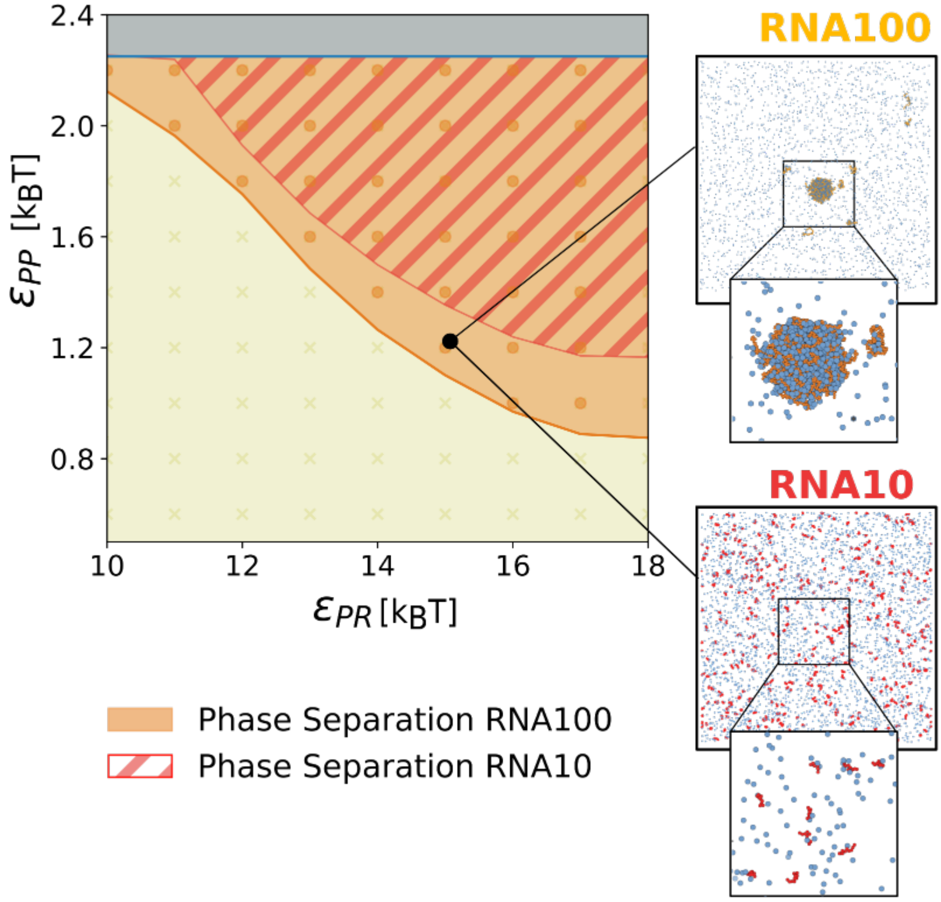


**Figure S4.** Phase diagram as a function of $\varepsilon_{PR}$ and $\varepsilon_{PP}$. Each point corresponds to an independent simulation. The orange region denotes parameter combinations for which condensates form in the presence of RNA chains of length 100, whereas the red region identifies conditions under which phase separation occurs even with RNA chains of length 10. Representative simulation snapshots are shown on the right for RNA100 and RNA10 using identical interaction parameters $\varepsilon_{PP} = 1.2\ k_BT$, $\varepsilon_{PR} = 15\ k_BT$ and RNA monomer concentrations. Under these conditions, only RNA100 induces phase separation, highlighting the critical role of RNA length in promoting demixing.

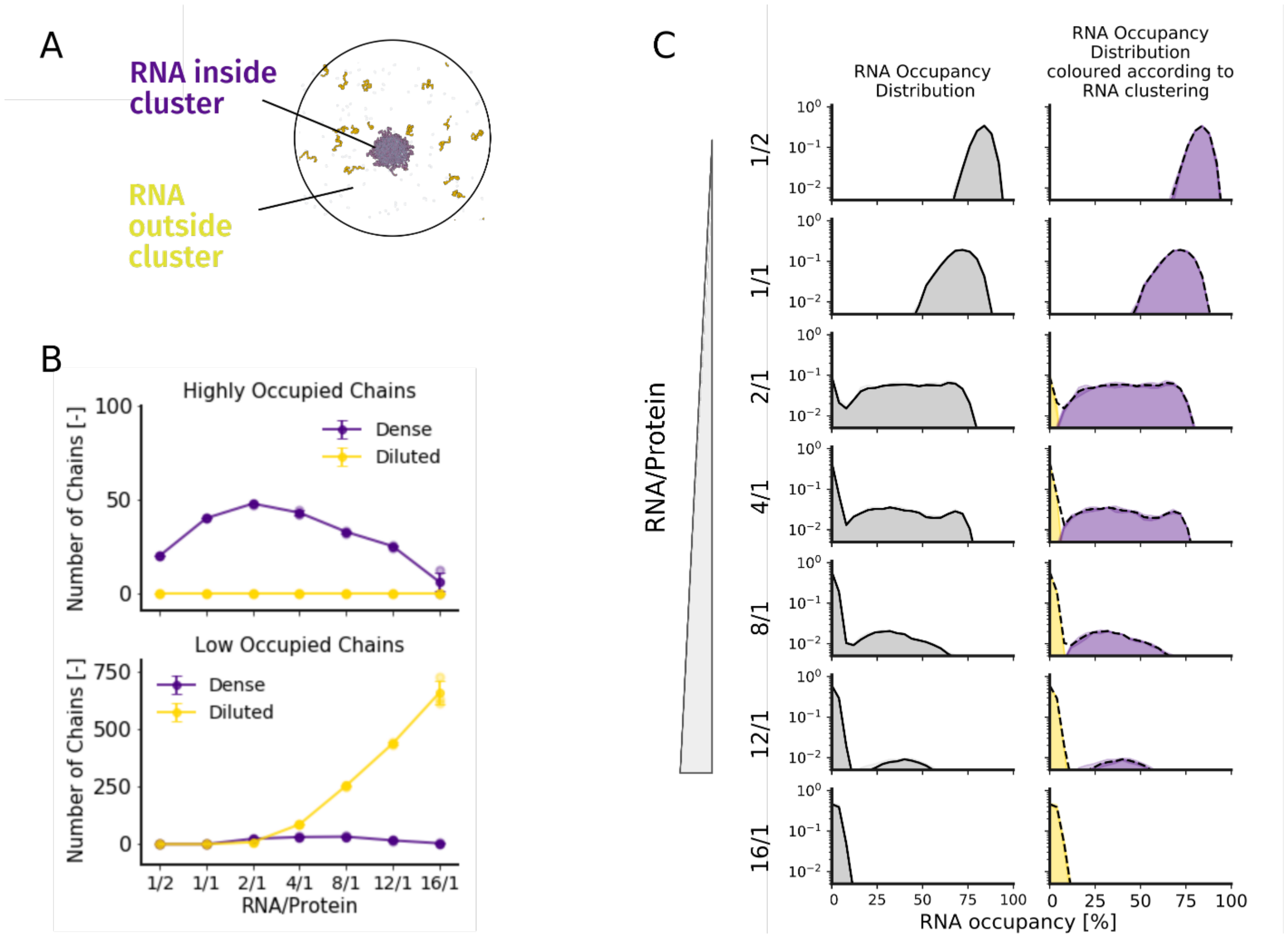


**Figure S5.** RNA occupancy and clustering. (A) Representative simulation snapshot in which RNA molecules are colored according to their clustering classification: purple indicates RNA molecules belonging to the condensate (dense phase), while yellow indicates RNA molecules located in the dilute phase. (B) Top: number of RNA chains with protein occupancy greater than 35% as a function of the RNA-protein ratio. Bottom: number of RNA chains with protein occupancy lower than 35% as a function of the RNA-protein ratio. In both panels, purple and yellow denote RNA molecules in the condensate and dilute phases, respectively. (C) Distribution of RNA occupancy as a function of the RNA-protein ratio. Left: occupancy distribution computed over all RNA molecules irrespective of phase localization (gray). Right: occupancy distributions computed separately for RNA molecules within the condensate (purple) and in the dilute phase (yellow).

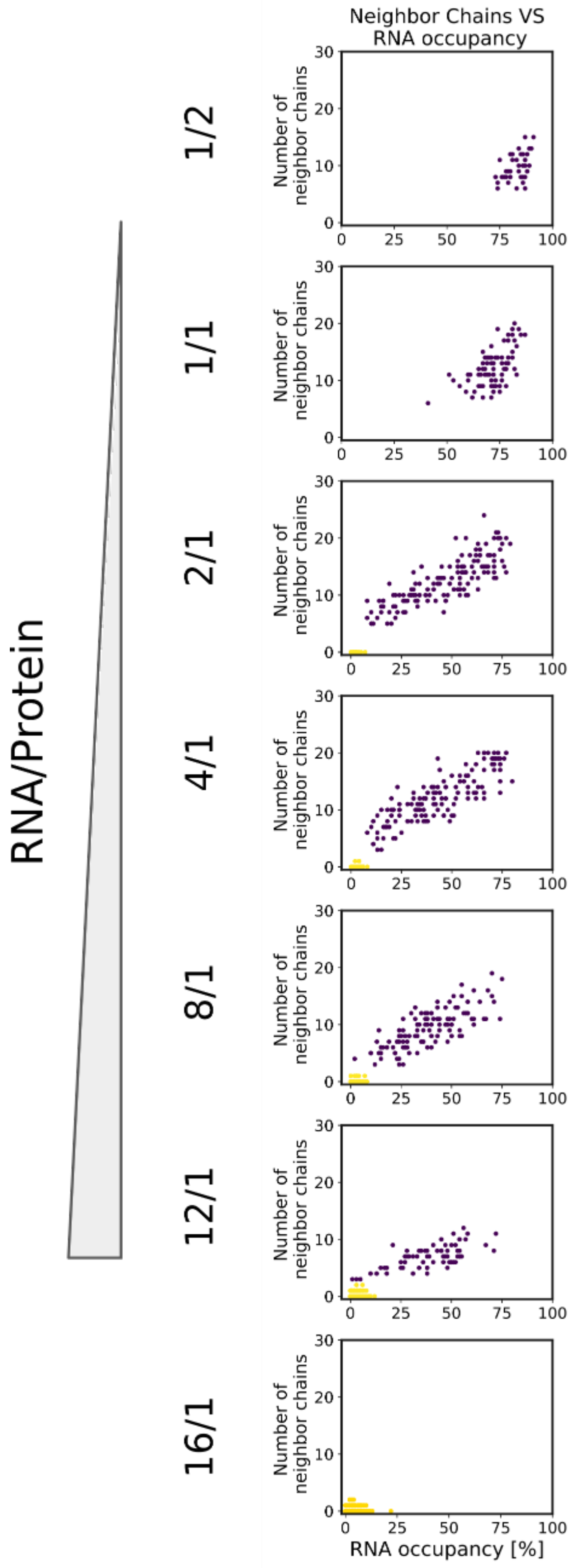


**Figure S6.** RNA clustering. Scatter plot of RNA chain occupancy versus the number of neighboring RNA chains at different RNA-protein ratios.

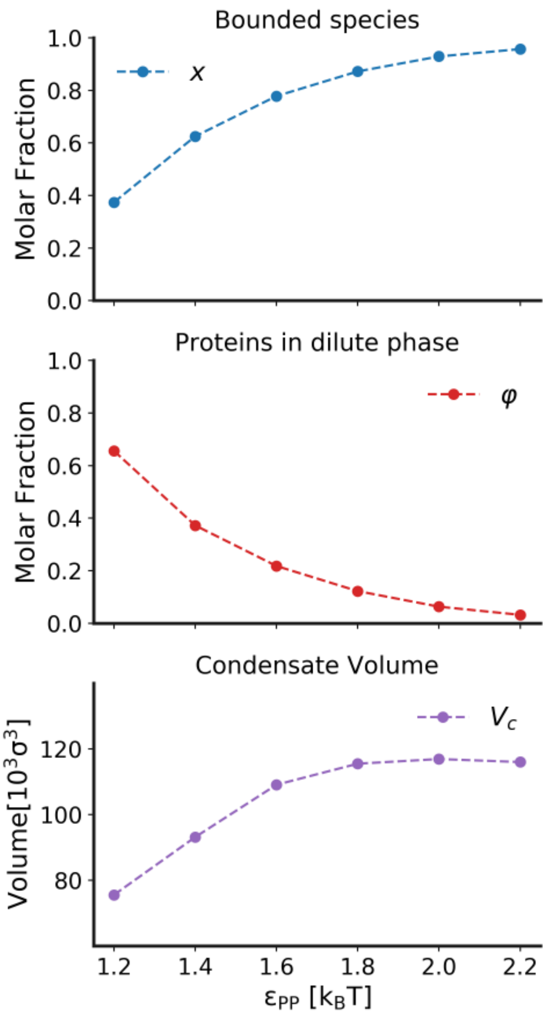


**Figure S7.** Dissociation constant within the condensate. The data shown correspond to the simulations reported in Figure 4 $\varepsilon_{PR} = 15\ k_BT$ and were used to calculate the condensate dissociation constant, $K_d^{\text{cond.}}$. For simulations exhibiting condensate formation ($\varepsilon_{PP} > 1.2\ k_BT$), the bound-species molar fraction, $x$ (blue, top panel), the molar fraction of proteins in the dilute phase, $\varphi$ (red, middle panel), and the condensate volume $V_c$ (purple, bottom panel) are reported as functions of $\varepsilon_{PP}$.
